# Generation of *KARRIKIN INSENSITIVE2* loss-of-function mutants in *Ceratopteris richardii* using a CRISPR/Cas9 system based on ribozyme-gRNA-ribozyme (RGR) technology

**DOI:** 10.64898/2026.04.22.720085

**Authors:** Ayano Wu, Yoshiya Seto, Junko Kyozuka, Yuki Hata

**Affiliations:** Graduate School of Life Sciences, Tohoku University, Katahira 2-1-1 Aoba-ku, Sendai, Japan; Laboratory of Plant Chemical Regulation, Department of Agricultural Chemistry, School of Agriculture, Meiji University, Kanagawa 214-8571, Japan; Deptartment of Agrotechnology & Food Sciences, Wageningen University, Wageningen, Stippeneng 2, 6708 WE, the Netherlands

**Keywords:** Gene editing, ferns, *Ceratopteris richardii*, KAI2, MAX2, SMXL

## Abstract

Plant hormones regulate almost every aspect of plant growth and development. KARRIKIN INSENSITIVE 2 (KAI2)-dependent signaling, which is thought to transduce signals derived from an unidentified ligand known as the KAI2 ligand (KL), regulates numerous traits, including seed germination in angiosperms and vegetative reproduction in bryophytes. The origin of KAI2 is believed to be ancient, and the evolution of its signaling pathways remains of significant interest. Ferns represent critical lineages for elucidating the evolution of land plant traits and growth mechanisms that enabled adaptation to terrestrial environments. Therefore, functional studies of key components of this pathway in ferns are essential for understanding the evolutionary trajectory of KAI2-dependent signaling during vascular plant diversification. However, experimental platforms for the CRISPR/Cas9 system, a powerful tool for investigating gene function, remain undeveloped in ferns. Here, we report an efficient CRISPR/Cas9 system based on ribozyme-gRNA-ribozyme (RGR) technology in the model fern, *Ceratopteris richardii* (*C. richardii*). We generated loss-of-function mutants of the *KAI2* ortholog in *C. richardii* (*CrKAI2*), as well as the signaling components *CrMAX2* and *CrSMXL*. We demonstrate that exogenous application of an artificial KL agonist increases the expression of KAI2-dependent signaling responsive genes in wild type plants; this response is abolished in *Crkai2* mutants. These findings indicate that KAI2-dependent signaling is conserved in *C. richardii*. Furthermore, this study proposes an efficient CRISPR/Cas9 method that will facilitate genetic studies in ferns.

## Introduction

Ferns, also known as monilophytes, represent the second-largest group in the plant kingdom (PPG I, 2016). They are evolutionarily significant lineages for understanding land plant evolution and offer unique biological resources (Yang et al., 2025b, Cao et al., 2017, Goswami et al., 2016, Shukla et al., 2016). Ferns propagate through spores, which germinate into free-living gametophytes (Fig. 1A) (Conway and Di Stilio, 2020). These gametophytes develop sexual organs, archegonia and antheridia, producing egg cells and sperm, respectively. The released sperm swim through water to reach the egg cell outside the gametophyte body, where fertilization occurs, and the resulting zygote develops into a sporophyte. During embryogenesis, the sporophyte receives nutrients from the gametophyte via the calyptra but becomes independent during post-embryonic development. The mature sporophyte exhibits complex morphology, including leaves, stems, roots, and vascular tissues. Upon maturation, the life cycle is completed with the formation of sporangia, which produce spores. Ferns and lycophytes form paraphyletic groups within the vascular plant lineage, with ferns identified as the sister group to seed plants (PPG I, 2016). The diversification of their ancestors involved several key evolutionary innovations, including the transition to a sporophyte-dominant life cycle, the development of leaves and roots, and the evolution of a vascular system (Jill Harrison, 2017, Harrison, 2017, Paul Kenrick, 1997, Graham et al., 2000). Thus, uncovering the molecular mechanisms underlying fern growth is critical for understanding how land plants evolved these complex traits. Furthermore, because fern lineages occupy an intermediate evolutionary position between bryophytes and seed plants, they are indispensable for elucidating how mechanisms in non-vascular plants were conserved or diversified in vascular plants (Plackett et al., 2015a).

**Figure 1.**
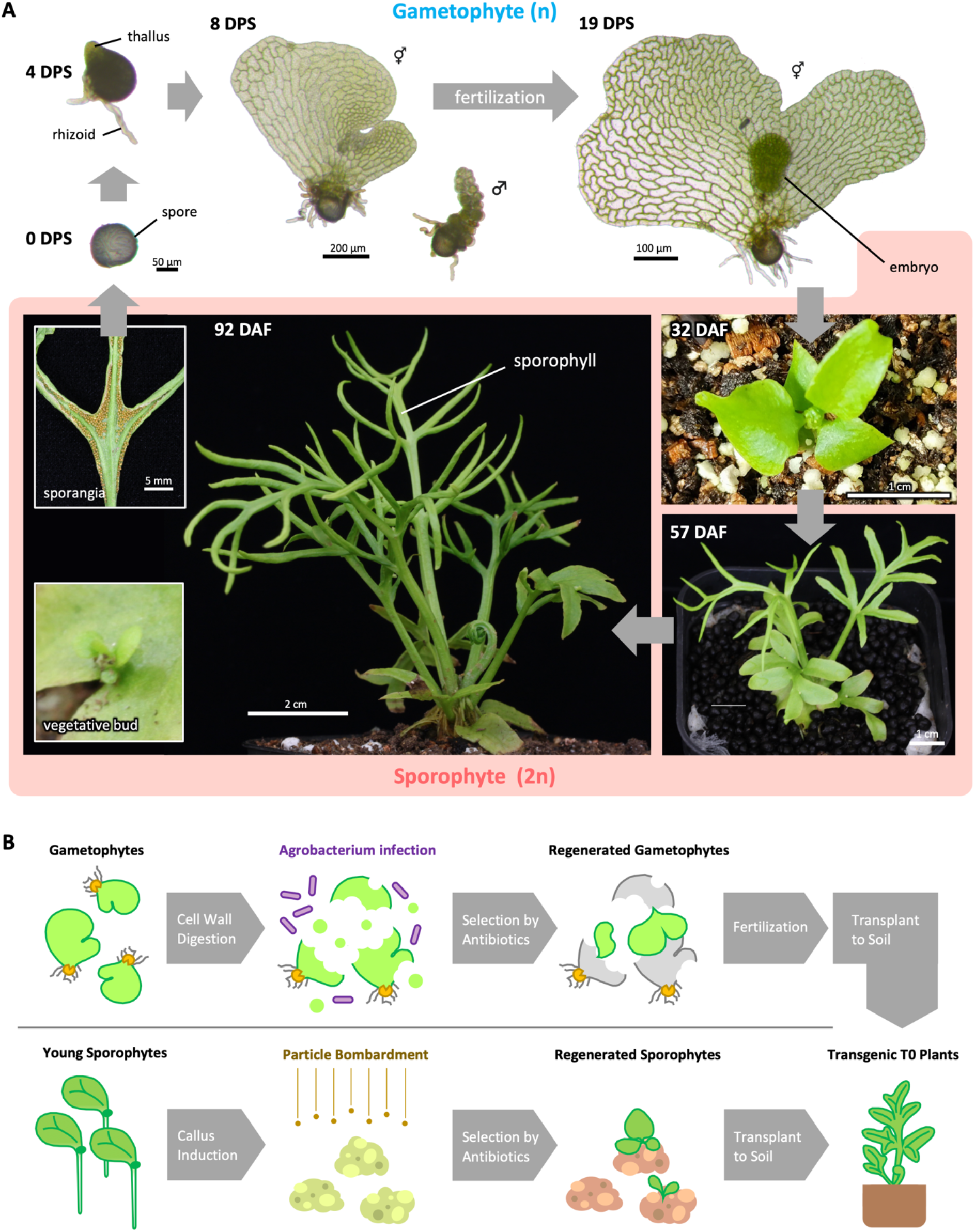
Life cycle and transformation of the fern *Ceratopteris richardii*. **(A)** Life cycle from 0 days after spore sowing (DPS) to 92 days after fertilization (DAF). The gametophyte phase starts from spore germination, producing rhizoids and a amall thallus. Gametophytes develop into either male (♂) or hermaphrodite (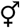) individuals. Male plants cease growing early and produce antheridia on their entire surface. Hermaphrodite plants form a concave notch meristem and continue growing and develop into a heart-shaped thallus. Archegonia are formed around the notch meristem, and antheridia are produced in the differentiated regions of the thallus. Sperms released from antheridia swim to archegonia, leading to fertilization. The sporophyte phase starts with this zygote formation. At first, the zygotes grow into embryos inside the parental gametophyte. They continue to grow and produce leaves and roots, becoming independent plant individuals. In the very initial stage of the sporophyte, the leaf shape is simple, but in the later stage, the shape of new leaves gradually becomes complex. Simultaneously, during the young sporophyte, they develop vegetative leaves with a wide width. However, when they are mature, sporophylls with a narrow width are produced, and sporangia are formed on the underside of a sporophyll. In some cases, vegetative buds are produced at an incision along the leaves. **(B)** Procedures of transformation. In the agrobacterium-mediated transformation (upper panels), the agrobacterium infects gametophytes following cell wall digestion to facilitate infection efficiency. After the transfer to a medium containing antibiotic, gametophytes harboring transgenes regenerate. Regenerated plants are fertilized to get transgenic T0 sporophytes. In the particle bombardment-mediated transformation (lower panel), calli induced from young sporophyte shoots are used for the bombardment. Transgenic T0 sporophytes are regenerated under medium containing antibiotic.

Despite the importance of fern research, the molecular mechanisms underlying fern growth remain poorly understood, largely due to their large genome sizes and the lack of suitable model systems (Plackett et al., 2015a, Ali et al., 2025). The average fern genome is approximately 12.3 billion bases (Gb), and ferns also exhibit high chromosome numbers (average, 40.5; maximum, 720), making genome sequencing particularly challenging (Ali et al., 2025). To date, genome resources are available only for twelve species (Marchant et al., 2022, Pelosi et al., 2025, Huang et al., 2022, Zhong et al., 2022, Song et al., 2025, Li et al., 2018, Rahmatpour et al., 2023, Qin et al., 2024, Xia et al., 2025, Fang et al., 2022, Shu et al., 2025). The lack of efficient transformation protocols has also been a major limitation. A few exceptions include *Pteris vittata* and *Ceratopteris thalictroides*, for which stable transformation via particle bombardment or *Agrobacterium tumefaciens* was reported in 2013; however, no genome sequences have been published for these species to date (Muthukumar et al., 2013). Subsequently, transformation protocols were developed for *Ceratopteris richardii* (*C. richardii*), commonly known as “C-fern,” which has been widely used as an educational model due to its rapid and easy growth (Kinosian and Wolf, 2022). The genome of *C. richardii* was sequenced at the chromosome scale in 2022, and studies employing genetic tools have recently begun to emerge (Kinosian and Wolf, 2022, Marchant et al., 2022). Although *C. richardii* is an aquatic fern, it shares key characteristics of leptosporangiate ferns, the largest taxon within monilophytes, including complex leaves (fronds) that develop from fiddleheads, homospory, and a heart-shaped gametophyte (Marchant et al., 2022). Four other aquatic ferns (*Azolla filiculoides, Azolla caroliana, Marsilea vestita*, and *Salvinia cucullata*) have also had their genome sequenced (Li et al., 2018, Song et al., 2025, Rahmatpour et al., 2023). However, these species exhibit specialized morphologies adapted to aquatic environments, and no transformation protocols have been reported for them (de Vries and de Vries, 2018, Babenko et al., 2019). As a result, *C. richardii* has emerged as the primary model of fern (Kinosian and Wolf, 2022). For genetic transformation, both particle bombardment in sporophytic calli and *Agrobacterium*-mediated transformation in gametophyte are available (Fig. 1B) (Plackett et al., 2014, Plackett et al., 2015b, Bui et al., 2015). To date, eight studies have used particle bombardment, while six have employed *Agrobacterium*-mediated transformation (Lai et al., 2026, Bui et al., 2017, Youngstrom et al., 2019, Youngstrom et al., 2022, Withers et al., 2023, Jiang et al., 2024, Renninger et al., 2025, Plackett et al., 2018, Geng et al., 2022, Xiang and Li, 2024, Geng et al., 2024, Yang et al., 2025a, McConnell et al., 2026, Schulz and Theißen, 2025).

Technologies of the clustered regularly interspaced short palindromic repeats (CRISPR)/CRISPR-associated nuclease 9 (Cas9) system have developed rapidly over the past two decades, making this approach indispensable for model plant research (Tuncel et al., 2025, Hassan et al., 2021). Recent innovations have enabled precise editing of DNA sequences across diverse plant species, targeting various genes and loci simultaneously (Molla et al., 2021, Zhang et al., 2021, Stuttmann et al., 2021). However, there are still difficulties in applying the CRISPR/Cas9 system across different species. Optimization of vector constructs for each plant species is an obstacle to the success of gene editing.

Application of CRISPR/Cas9 in *C. richardii* has also been reported in several recent studies; however, there is still considerable room for improvement (Jiang et al., 2024, Xiang and Li, 2024, Schulz and Theißen, 2025, Lai et al., 2026). One critical consideration is the strategy for guide RNA (gRNA) expression. gRNA must be actively transcribed at the appropriate timing following transformation and retained within the nucleus to form a functional complex with Cas9. Because transcripts produced by RNA polymerase II are exported from the nucleus, gRNA expression is commonly driven by RNA polymerase III using U3 or U6 promoters. However, these promoters are often species-specific and may not function efficiently across distantly related plant lineages. Furthermore, U3 and U6 promoters are generally constitutively active, limiting the ability to fine-tune expression levels and timing to improve editing efficiency (Hassan et al., 2021, Gao and Zhao, 2014). One promising approach to overcome these limitations is ribozyme-gRNA-ribozyme (RGR) technology (Gao and Zhao, 2014). In this system, the transcribed gRNA includes self-cleaving ribozymes that remove flanking sequences through autocatalytic reactions, preventing nuclear export signals from being retained. As a result, the RGR construct can be expressed using a wide range of polymerases and promoters, offering greater flexibility and overcoming constraints associated with conventional gRNA expression systems.

KARRIKIN INSENSITIVE 2 (KAI2) -dependent signaling is a conserved molecular pathway in land plants that is stimulated by an unidentified plant hormone, referred to as “KL” (Waters and Smith, 2013, Bythell-Douglas et al., 2017). Upon the ligand perception, the receptor KAI2 forms a complex with the F-box protein MORE AXILLARY GROWTH 2 (MAX2) and the transcriptional repressor SUPPRESSOR OF MAX2 1-Like (SMXL) (Stanga et al., 2013, Wang et al., 2015, Stanga et al., 2016). SMXL is subsequently degraded via the proteasome pathway, leading to activation of downstream gene expression. Although these signaling components are highly conserved across land plants, their functions have diverged during evolution. In angiosperms, KAI2-dependent signaling promotes seed germination (Nelson et al., 2012). In contrast, in early-diverging land plants, KAI2-dependent signaling regulates vegetative propagation processes, such as gemma production in *Marchantia polymorpha* (*M. polymorpha*), and gametophore formation in *Physcomitrium patens* (*P. patens*) (Komatsu et al., 2023, Komatsu et al., 2025, Luo et al., 2025, Guillory et al., 2024, Lopez-Obando et al., 2021). Furthermore, gene duplication of KAI2 in the common ancestor of seed plants led to the evolution of DWARF14 (D14), a strigolactone receptor that regulates diverse developmental processes, including shoot branching (Bythell-Douglas et al., 2017). In addition, root-parasitic plants have evolved the ability to perceive host-derived strigolactones through receptors derived from KAI2 duplication and diversification (Conn et al., 2015). In angiosperms, KAI2 detects smoke-derived karrikins and promotes seed germination (Nelson et al., 2012). However, the evolutionary trajectory of the KAI2-dependent signaling and the origin of SL signaling in early vascular plants remain poorly understood. *C. richardii* represents an ideal model to address these questions. In this study, we developed an efficient CRISPR/Cas9 system in *C. richardii* targeting genes involved in the KAI2-dependent signaling.

## Results

### A construct containing *Marchantia* promoters failed to induce editing in *C. richardii*

First, we assessed KAI2 orthologs are conserved across monilophytes, including *C. richardii*. We performed BLAST searches using genome data from several fern species and constructed a phylogenetic tree. All analyzed fern species possess conserved proteins belonging to the eu-KAI2 superclade (KAI2 orthologs), supporting the high conservation of KAI2 proteins in monilophytes (Fig. 2A). In *C. richardii*, a single-copy protein (Ceric.28G021700) clusters within the eu-KAI2 clade. Its amino acid sequence shows high similarity to both AtKAI2 and MpKAI2A, including the conserved catalytic triad, key residues required for ligand perception and signaling (Fig. 2B) (Hamiaux et al., 2012). Based on these features, we identified this protein as a KAI2 ortholog and designated it CrKAI2.

**Figure 2.**
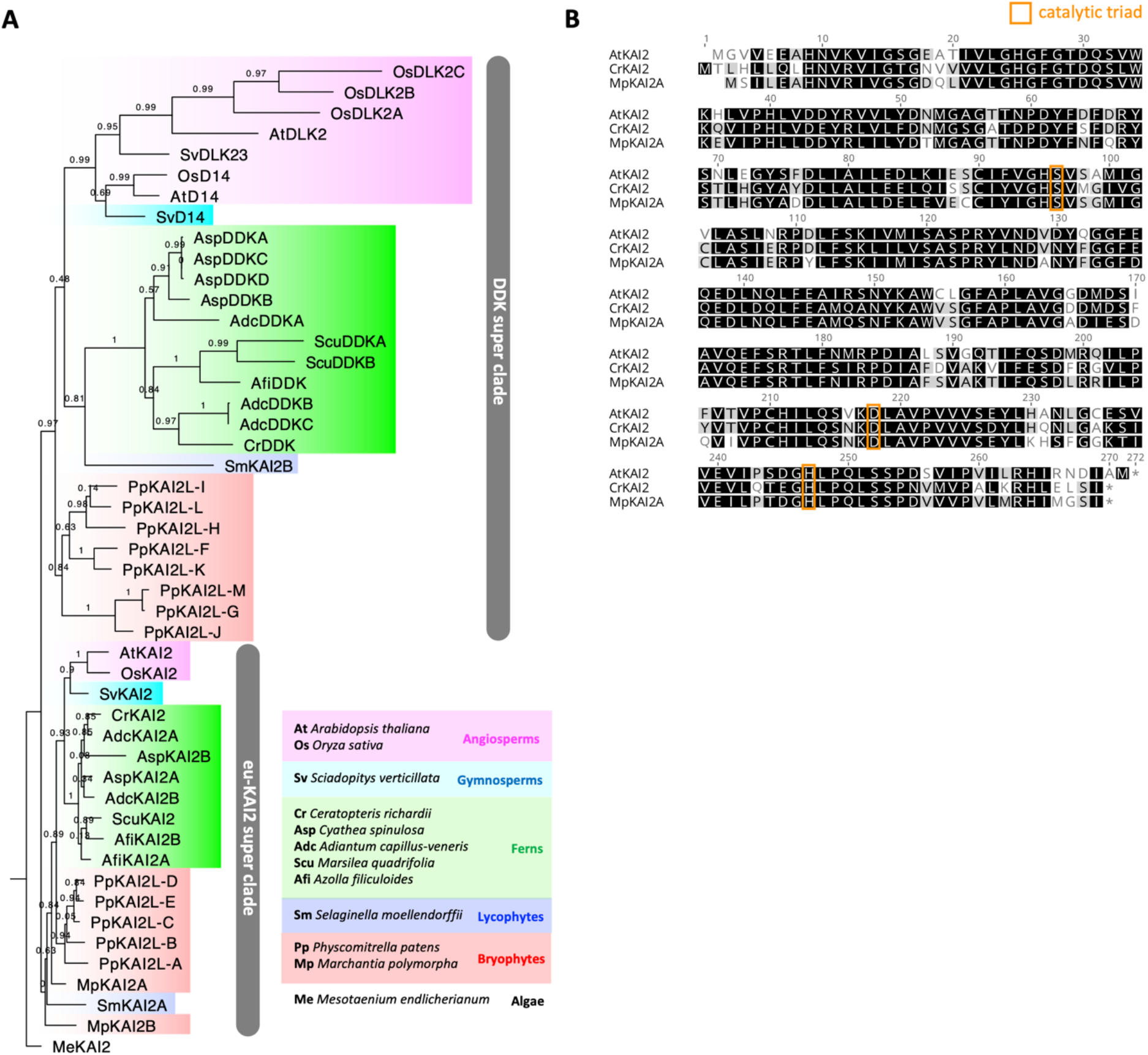
KAI2 orthologs are conserved in Ferns. **(A)** Phylogenetic tree of KAI2 and D14 family proteins. **(B)** Alignment of amino acid sequences of KAI2 proteins from *A. thaliana, C. richardii*, and *M. polymorpha*. Residues comprising the catalytic triad are highlighted with orange rectangles.

Next, we evaluated whether CRISPR/Cas9 constructs developed for *M. polymorpha* (pMpGE010 and pMpGE_En03) are applicable to *C. richardii* (Fig. 3A) (Sugano et al., 2018). In this system, the MpU6 promoter (an endogenous promoter from *M. polymorpha*) and the gRNA scaffold are cloned into the pMpGE_En03 vector, while Cas9-NLS driven by the MpEF promoter is contained in pMpGE010. To target a specific genome region, the gRNA sequence in pMpGE_En03 is modified, and the resulting gRNA expression cassette is integrated into pMpGE010 via an LR reaction.

**Figure 3.**
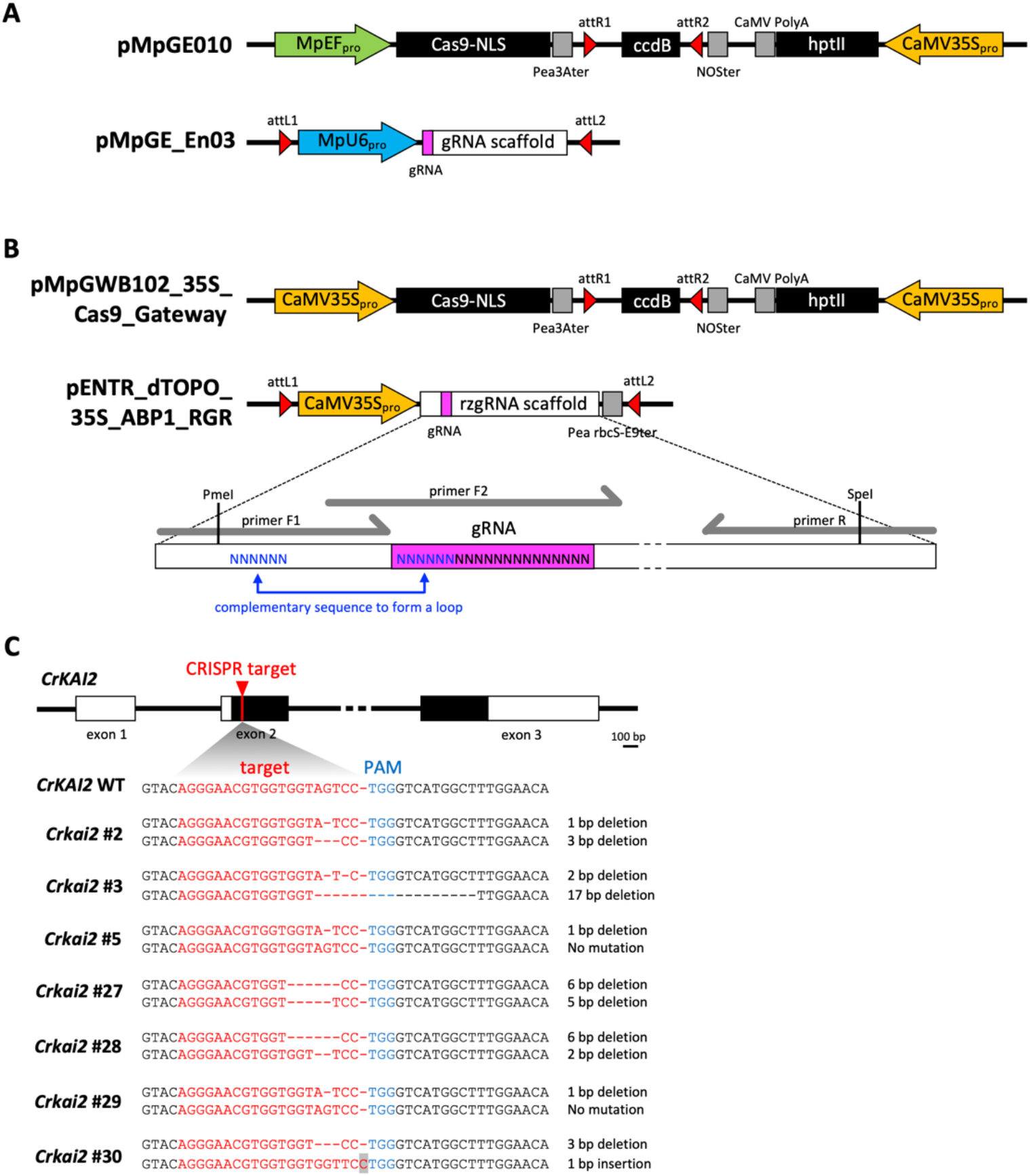
The RGR CRISPR/Cas9 system efficiently introduces editing in *CrKAI2*. **(A)** Schematic diagrams of gene editing constructs, diverted from *M. polymorpha* to *C. richardii*. **(B)** Constructs developed in this study using the RGR-based technology. Arrows filled with a color, boxes filled with black, and boxes filled with gray indicate promoters, protein-coding sequences, and terminators. gRNA (rzgRNA) scaffolds and gRNA sequences were shown by white boxes and magenta color. Primer F1, F2, and R indicates positions of primers used for over extension PCR to modify the gRNA sequence in the rzgRNA scaffold. **(C)** Gene structure of *CrKAI2* and introduced mutation patterns at the CRISPR/Cas9 target site. White and black boxes indicate exon and protein-coding sequences, respectively.

We selected a target sequence within the second exon of *CrKAI2*, located just downstream of the start codon. A CRISPR/Cas9 vector for *CrKAI2* editing was constructed based on the Marchantia system (pMpGE010) and introduced into calli via particle bombardment. The transformed calli yielded 29 regenerated plants, of which 23 were stably transformed (Table 1). Integration of the construct into the genome was confirmed by genotyping; however, no plants exhibited gene editing at the target site.

**Table 1.** Number of plant individuals after the transformation.

| Construct used | target | regenerated | stably transformed | edited |
| --- | --- | --- | --- | --- |
| MpEFpro:Cas9 ;<br>MpU6pro:CrKAI2gRNA | *CrKAI2* | 29 | 23 (79%) | 0 (0%) |
|  | *CrMAX2* | 5 | 4 (80%) | 0 (0%) |
|  | *CrSMXL* | 11 | 9 (82%) | 0 (0%) |
| 35Spro:Cas9 ;<br>35Spro:CrKAI2RGR | *CrKAI2* | 30 | 21 (70%) | 10 (33%) |
|  | *CrMAX2* | 32 | 17 (53%) | 2 (6%) |
|  | *CrSMXL* | 34 | 24 (71%) | 3 (9%) |

### A CRISPR/Cas9 system using the RGR-based strategy successfully induced genome editing

Although the U6 promoter can function across land plant species, its applicability between distantly related species is limited. Therefore, in a second approach, we developed a CRISPR/Cas9 system based on the ribozyme-gRNA-eibozyme (RGR) strategy (Fig. 3B) (Gao and Zhao, 2014). To facilitate modification of the gRNA sequence, we adopted a vector design similar to that used in Marchantia. The vector pENTR_dTOPO_35S_ABP1_RGR contains a ribozyme-gRNA (rzgRNA) scaffold, while pMpGWB102_35S_Cas9_Gateway carries the Cas9 expression cassette. The rzgRNA sequence can be easily modified by PCR using primers encoding the desired target sequence. The modified rzgRNA expression cassette is then introduced into the pMpGWB102_35S_Cas9_Gateway via LR recombination reaction. In this system, the rzgRNA scaffold, Cas9-NLS, and the antibiotic selection marker are all driven by the CaMV35S promoter, which is currently reliable constitutive promoter in *C. richardii* (Plackett et al., 2015b).

Calli transformed with the RGR-based CRISPR/Cas9 vector (pMpGWB102_35S_Cas9_Gateway) regenerated 30 plants, of which 21 were confirmed to carry the transgene (Table 1). In contrast to the results obtained with pMpGE010, we identified 10 plants (33%) harboring edits at the target site. A range of editing patterns was observed, including both insertions and deletions (Fig. 3C). Several mutations resulted in frameshift and/or premature termination of the coding sequence. Among the seven mutant plants shown in Fig. 3C, two (lines #5 and #9) are heterozygous, while the remaining five (lines #2, #3, #27, #28 and #30) are compound heterozygous, carrying distinct mutations in both alleles. In line 3, mutations in both alleles cause a frameshift, leading to loss of *CrKAI2* function. These findings suggest that null mutants in *CrKAI2* are unlikely to be lethal.

### *Crkai2* mutants are insensitive to the treatment of (−)-GR24 treatment, a synthetic agonist of the KAI2-dependent signaling

To determine whether the introduced mutation in *CrKAI2* disrupts its function, we generated T1 plants homozygous for the mutation through self-crossing. Four T0 lines (#2, #28, #29, and #30) were used (Fig. 3C). After spores were harvested from each line, they were germinated on medium and self-crossed, and the genotypes of the resulting T1 sporophytes were determined. We successfully identified at least two T1 plants carrying homozygous mutations from each T0 line. T1 plants carrying homozygous single-base deletion were obtained from #2 and #29, whereas those with a homozygous two-base deletion were obtained from #28 and #30.

KAI2 functions as a receptor for KAI2 ligands; therefore, we hypothesized that *Crkai2* mutants would be insensitive to (−)-GR24, a synthetic agonist of this signaling pathway. To test this, we assessed the sensitivity of homozygous T1 plants to (−)-GR24 using #2-11 and #28-14 as representative homozygous mutants.

In angiosperms and bryophytes, (−)-GR24 stimulate KAI2-dependent signaling, thereby upregulating KL-responsive genes such as *KAR-UP F-box* (*KUF*), *SMXL*, and *DWARF14-LIKE2* (*DLK2*) (Stanga et al., 2013, Mizuno et al., 2021, Lopez-Obando et al., 2021, Sepulveda et al., 2022). BLAST searches and phylogenetic analyses revealed single copy orthologs of the *SMXL* (*CrSMXL*), *DLK2* (*CrDDK*), and *KUF* (*CrKUF*) in *C. richardii*, respectively (Fig. 2A and S1A). These three genes were used as markers. In wild-type (WT) plants, expression levels of *CrKUF* and *CrDDK* increased significantly following (−)-GR24 treatment (Fig. 4). Expression of *CrSMXL* was also slightly upregulated, although the change was not statistically significant. In contrast, in *Crkai2* mutants, baseline expression levels of all three marker genes were substantially lower than in WT under the control condition and their expression remained unchanged after (−)-GR24 treatment. These results indicate that *Crkai2* mutants are insensitive to both endogenous ligands and (−)-GR24.

**Figure 4.**
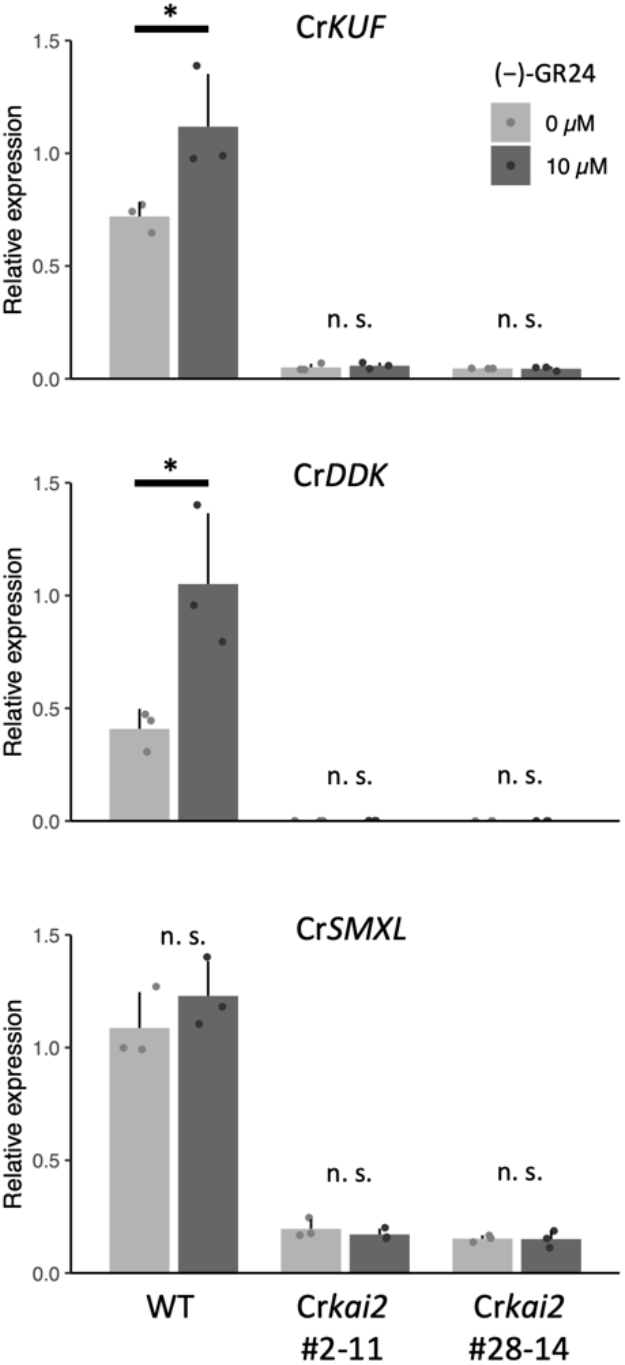
*Crkai2* mutants are insensitive to (−)-GR24 treatment. Expression levels of *CrKUF, CrDDK*, and *CrSMXL* in WT and *Crkai2* loss-of-function mutants (#2-11 and #28-14), measured by qRT-PCR. Statistical significance was examined by Student’s *t* test (*n* = 3, \**P* < 0.05).

### The RGR-based CRISPR/Cas9 system is efficient across multiple targets in *C. richardii*

We next evaluated whether the RGR-based CRISPR/Cas9 system could be applied to additional target genes. Guide RNA (gRNA) target sites were designed within the coding region of *CrSMXL* and *CrMAX2*, the latter being a single ortholog of *MAX2* found in the *C. richardii* genome (Fig. S1B). As in the CrKAI2 experiments, two constructs were generated for each target: one based on pMpGE010 and another using the RGR CRISPR/Cas9 vector. These constructs were introduced into *C. richardii* independently. In experiments using pMpGE010, no plants with edits at the target sites were identified (Table 1). In contrast, gene editing at the target loci was confirmed in multiple plants transformed with the RGR CRISPR/Cas9 vector (Fig. S1C and S1D). We found three plants (9%) harboring edits at the target site for *CrSMXL*, and two plants (6%) for *CrMAX2* (Table 1). Unlike T0 plants of *Crkai2* mutants, all mutations detected in both *Crsmxl* and *Crmax2* T0 plants were heterozygous. These findings further demonstrate the robust efficiency of the RGR-based CRISPR/Cas9 system across multiple targets in *C. richardii*.

## Discussion

We found that the RGR-based CRISPR/Cas9 system functions efficiently for gene editing in *C. richardii*. The observed insensitivity of the mutants to (−)-GR24 treatment strongly supports the successful disruption of *CrKAI2* gene function through genome editing. In contrast, the CRISPR/Cas9 system based on pMpGE010 did not produce any edits, despite generating a comparable number of regenerated plants as the RGR vector. Given that bryophytes and vascular plants, including monilophytes, diverged approximately 500 million years ago, it is likely that the MpU6 and/or MpEF1α promoters in pMpGE010 are not fully compatible with *C. richardii* (Harris et al., 2022).

Although previous studies have demonstrated that CRISPR/Cas9-mediated gene editing is feasible in *C. richardii*, our method offers several advantages (Jiang et al., 2024, Xiang and Li, 2024, Lai et al., 2026). First, we used the CaMV35S promoter to express rzgRNA. CaMV 35S is known to be highly active during callus growth and regeneration from calli (Plackett et al., 2015b). Therefore, our approach is broadly effective across various target genes. Successful editing of the three different genes further supports the robustness of target selection. In previous studies, gRNA was expressed under an endogenous U3 or U6 promoter; however, other U3 and U6 promoters were not evaluated (Xiang and Li, 2024, Lai et al., 2026). Therefore, it remains unclear whether the promoter used in these studies shows enough expression level for efficient editing in other targets. Another study employed CRISPR/Cas9 with gRNA expression under an endogenous ACTIN promoter (Jiang et al., 2024). However, this approach may reduce editing efficiency, as gRNA transcribed by RNA polymerase II could be exported from the nucleus. Second, our experimental design is simpler than those reported previously. The tRNA-based system used in earlier studies requires relatively complex construct assembly (Xiang and Li, 2024). Another study reported the establishment of transgenic lines that constitutively express Cas9 (Schulz and Theißen, 2025). These lines may serve as useful platforms for introducing gRNA-encoding constructs in the future, enabling more efficient genome editing. While this strategy may facilitate gene knockout in a WT background, it has limitations in certain transgenic context, such as reporter lines for gene expression. In contrast, our method employs a straightforward, single-vector system for callus transforming, offering a simple and efficient approach. This design is likely facilitating future genetics studies in *C. richardii*.

This study represents the first application of RGR-based CRISPR/Cas9 technology outside of angiosperms. Because the CaMV35S is widely used across diverse plant species, our constructs have the potential to be applied to species in which gene editing has not yet been reported, including ferns, bryophytes, and algae. Even if the CaMV35S promoter is not effective in certain species, the RGR-based CRISPR/Cas9 system does not impose strict limitations on promoter selection, allowing for straightforward optimization. Furthermore, tissue-specific gene knockout in ferns may be achieved by replacing the CaMV35S promoter with a tissue-specific promoter, as previously demonstrated in *Arabidopsis thaliana* (Gao et al., 2015).

Both the insensitivity to (-)-GR24 treatment and the reduced baseline expression of (-)-GR24-responsive genes in *Crkai2* mutants clearly demonstrate the conserved function of CrKAI2 as a KL receptor. Furthermore, the responsiveness of *SMXL, KUF*, and *DDK* gene expression to KL signaling is shared across land plants, including ferns, suggesting strong conservation of the KAI2-dependent signaling pathway (Stanga et al., 2013, Mizuno et al., 2021, Lopez-Obando et al., 2021, Sepulveda et al., 2022). Although the biological significances of these conserved responses remain unclear, it has been hypothesized that KUF inhibits KL biosynthesis, forming a negative feedback loop in *A. thaliana* (Sepulveda et al., 2022). Because SMXL proteins act as repressors of the signaling pathway, their upregulation likely also contributes to negative feedback regulation. Our results suggest conserved mechanisms that fine-tune KAI2-dependent signaling in *C. richardii*, implying an important role for this pathway in fern growth. The mutants generated in this study are expected to be valuable for uncovering the function of KAI2-dependent signaling in ferns and for understanding the evolutionary trajectory of this pathway across land plants.

Another intriguing question concerns the origin of a D14, the strigolactone receptor. D14 is thought to have evolved from KAI2 paralogs in the common ancestor of seed plants. *Marchantia* possesses KAI2 but not D14, and perceives only KL-mimic compounds (Kodama et al., 2022). *Physcomitrim patens* contains multiple KAI2 paralogs resulting from gene duplication; some of these can recognize strigolactones, although this ability is likely a derived trait in mosses (Lopez-Obando et al., 2021). In contrast, gymnosperms and angiosperms possess both KAI2 and D14 and perceive KL-mimic compounds and strigolactone, respectively (Kodama et al., 2023). This study demonstrates that a KL-mimic compound is perceived by a single CrKAI2 in *C. richardii*. It would be informative to test whether CrKAI2 can also recognize strigolactones, which would provide further insights into the evolutionary origin of strigolactone signaling.

## Materials and Methods

### Plant materials and growth conditions

All experiments were performed using *Ceratopteris richardii* strain Hn-n (Warne and Hickok, 1987). Plants were cultured and grown as described previously (Plackett et al., 2015b). In brief, spores were surface sterilized by soaking in 10% hypochlorous acid for 10 minutes, followed by five washes with sterilized water. The sterilized spores were incubated in the dark at room temperature for 3 days before transferring to C-fern medium. Gametophores and young sporophyte were cultured in 9 cm dishes overlaid with C-fern medium containing 1% agar (Nacalai Tesque) under a 16 hours light/8 hours dark cycle at 28°C. Adult sporophytes were transferred to soil-filled pots and grown in a greenhouse. To maintain humidity, pots were placed in transparent plastic containers with water.

### Preparation of (−)-GR24

*rac*-GR24 was prepared according to the procedure in a previous report (Chen et al., 2021). Each enantiomer of GR24 was then optically separated using a chiral column (CHIRAL PAK IC, 10 mm×250 mm, 100% MeOH, 2.0 ml/min) to afford (−)-GR24 at 17.2 min.

### Vector construction

For the construction of pENTR_dTOPO_35S_CrKAI2_RGR, pENTR_dTOPO_35S_CrMAX2_RGR, and pENTR_dTOPO_35S_CrSMXL_RGR, DNA fragments containing the CaMV35S promoter and rzgRNA scaffold were amplified by PCR using pHDE_35S_ABP1_RGR_AP1_Cas9 as a template and cloned into pENTR/d-TOPO (Gao et al., 2015). The rzgRNA sequence was modified for each target using overlap extension PCR. The pHDE_35S_ABP1_RGR_AP1_Cas9 vector was digested with PmeI and SpeI to remove the original rzgRNA sequence. The modified PCR products were then introduced into the digested vector using SLiCE reaction (Zhang et al., 2012).

For the construction of pMpGWB102_35S_Cas9_Gateway, a DNA fragment containing Cas9-NLS and the Pea3A terminator was amplified from pMpGE_010 and subcloned into pENTR/d-TOPO, followed by transfer to pMpGWB102 via LR recombination (Sugano et al., 2018). The Gateway cassette, amplified from pMpGWB303, was introduced into the Aor51HI restriction site. To generate the destination vector for plant transformation, the fragment containing the CaMV35S promoter and rzgRNA scaffold was introduced via LR recombination. For the construction of pMpGE010 targeting CrKAI2, CrMAX2, or CrSMXL, pMpGE_En03 was linearized with BsaI, and oligonucleotides encoding the gRNA sequence were ligated into the BsaI site. The modified gRNA scaffold and MpU6 promoter in pMpGE_En03 were then transferred into pMpGE010 via LR recombination. All primers used for vector construction are listed in Table S1.

### Transformation and genotyping

Transformation of *C. richardii* was performed as described by Plackett et al. 2015, with minor modifications (Plackett et al., 2015b). Young sporophytes with a single leaf were transferred to Murashige and Skoog (MS) medium containing 0.5 µM 6-benzylaminopurine (BAP) to induce callus formation. After two weeks under standard growth conditions, calli formed at the shoot tips were collected and propagated on MS medium supplemented with BAS. Prior to transformation, calli were transferred to MS medium containing 5 µM kinetin (KT) and cultured for 2 days. Transformation was carried out using a PDS-1000/He particle delivery system (Bio-Rad) with 1.6 µm gold particles and rupture disks at 1100 psi. The following day, bombarded calli were transferred to MS medium containing 5 µM KT and 40 mg/L hygromycin B and cultured for two weeks. Regenerated calli were maintained on selection medium, and regenerated sporophytes were transferred to C-fern medium. Genomic DNA was extracted from approximately 7th or 8th leaf. Leaves were homogenized using a Multi-Beads Shocker (Yasui Kikai) and DNA was extracted using the CTAB method. Target regions were amplified by PCR and analyzed by Sanger sequencing. All primers used for genotyping are listed in Table S1.

### Gene expression analysis

Young sporophytes with 10-12 fully expanded leaves were used for gene expression analyses by quantitative RT-PCR. The second, third, and forth youngest leaves from each plant were collected and pooled for treatment with 10 µM (-)-GR24 or a control solution containing an equivalent amount of acetone. Treatment was performed by soaking leaves in the solution and incubating them for 3 hours under standard growth conditions, followed by 10 minutes of vacuum-infiltration. Leaf tissues were then frozen in liquid nitrogen and ground using a pre-chilled mortar. Total RNA was extracted using the RNeasy Plant Mini Kit (Qiagen). Then, cDNA was synthesized using SuperScript VILO cDNA Synthesis Kit (Invitrogen). Quantitative PCR was performed using a CFX Duet system (Bio-Rad) with KOD SYBR qPCR mix (Toyobo), synthesized cDNA, and primers listed in Table S1.

### Phylogenetic analysis

Amino acid sequences of D14, KAI2, MAX2, and SMAX1 proteins in *A. thaliana* were used as queries for BLAST search against Phytozome v13 and FernBase. Retrieved sequences were aligned using the MUSCLE algorithm, and gaps were manually curated using CLC sequence viewer 8. Phylogenetic trees were constructed using the maximum likelihood method implemented in PhyML

## Data Availability

The data underlying this article are available in the article and in its online supplementary material.

## Acknowledgements

We thank Mitsuyasu Hasebe (NIBB, Japan) for providing wild-type spores of *C. richardii*; Andrew Plackett Yunde Zhao (University of California San Diego, United States) for providing pHDE_35S_ABP1_RGR_AP1_Cas9.

## References

Ali, Z., Tan, Q.W., Lim, P.K., Chen, H., Pfeifer, L., Julca, I., et al. (2025) Comparative transcriptomics in ferns reveals key innovations and divergent evolution of the secondary cell walls. Nature Plants. 11: 1028–1048.

Babenko, L., Vasheka, O., Shcherbatiuk, M., Romanenko, P., Voytenko, L., and Kosakivska, I. (2019) Biometric characteristics and surface microstructure of vegetative and reproductive organs of heterosporous water fern Salvinia natans. Flora. 252: 44–50.

Bui, L.T., Cordle, A.R., Irish, E.E., and Cheng, C.-L. (2015) Transient and stable transformation of Ceratopteris richardii gametophytes. BMC Research Notes. 8: 214.

Bui, L.T., Pandzic, D., Youngstrom, C.E., Wallace, S., Irish, E.E., Szövényi, P., et al. (2017) A fern AINTEGUMENTA gene mirrors BABY BOOM in promoting apogamy in Ceratopteris richardii. The Plant Journal. 90: 122–132.

Bythell-Douglas, R., Rothfels, C.J., Stevenson, D.W.D., Graham, S.W., Wong, G.K.-S., Nelson, D.C., et al. (2017) Evolution of strigolactone receptors by gradual neo-functionalization of KAI2 paralogues. BMC Biology. 15: 52.

Cao, H., Chai, T.-T., Wang, X., Morais-Braga, M.F.B., Yang, J.-H., Wong, F.-C., et al. (2017) Phytochemicals from fern species: potential for medicine applications. Phytochemistry Reviews. 16: 379–440.

Chen, Y., Kuang, Y., Shi, L., Wang, X., Fu, H., Yang, S., et al. (2021) Synthesis and Evaluation of New Halogenated GR24 Analogs as Germination Promotors for Orobanche cumana. Frontiers in Plant Science. 12: 725949.

Conn, C.E., Bythell-Douglas, R., Neumann, D., Yoshida, S., Whittington, B., Westwood, J.H., et al. (2015) Convergent evolution of strigolactone perception enabled host detection in parasitic plants. Science. 349: 540–543.

Conway, S.J., and Di Stilio, V.S. (2019) An ontogenetic framework for functional studies in the model fern Ceratopteris richardii. Developmental Biology. 457: 20–29.

De Vries, S., and De Vries, J. (2018) Azolla: A Model System for Symbiotic Nitrogen Fixation and Evolutionary Developmental Biology. In Current Advances in Fern Research. pp. 21–46.

Fang, Y., Qin, X., Liao, Q., Du, R., Luo, X., Zhou, Q., et al. (2022) The genome of homosporous maidenhair fern sheds light on the euphyllophyte evolution and defences. Nature Plants. 8: 1024–1037.

Gao, Y., Zhang, Y., Zhang, D., Dai, X., Estelle, M., and Zhao, Y. (2015) Auxin binding protein 1 (ABP1) is not required for either auxin signaling or Arabidopsis development. Proceedings of the National Academy of Sciences. 112: 2275–2280.

Gao, Y., and Zhao, Y. (2013) Self-processing of ribozyme-flanked RNAs into guide RNAs in vitro and in vivo for CRISPR-mediated genome editing. Journal of Integrative Plant Biology. 56: 343–349.

Geng, Y., Xie, C., Yan, A., Yang, X., Lai, D.N., Liu, X., et al. (2024) A conserved GRAS-domain transcriptional regulator links meristem indeterminacy to sex determination in Ceratopteris gametophytes. Current Biology. 34: 3454-3472.e7.

Geng, Y., Yan, A., and Zhou, Y. (2022) Positional cues and cell division dynamics drive meristem development and archegonium formation in Ceratopteris gametophytes. Communications Biology. 5: 650.

Goswami, H.K., Sen, K., and Mukhopadhyay, R. (2016) Pteridophytes: evolutionary boon as medicinal plants. Plant Genetic Resources. 14: 328–355.

Graham, L.E., Cook, M.E., and Busse, J.S. (2000) The origin of plants: Body plan changes contributing to a major evolutionary radiation. Proceedings of the National Academy of Sciences. 97: 4535–4540.

Guillory, A., Lopez-Obando, M., Bouchenine, K., Bris, P.L., Lécureuil, A., Pillot, J.-P., et al. (2024) SUPPRESSOR OF MAX2 1-LIKE (SMXL) homologs are MAX2-dependent repressors of Physcomitrium patens growth. The Plant Cell. 36: 1655–1672.

Guillory, A., Lopez-Obando, M., Bouchenine, K., Lambret, L., Bris, P.L., Lécureuil, A., et al. (2023) Physcomitrium patens SMXL homologs are PpMAX2-dependent negative regulators of growth. bioRxiv (Cold Spring Harbor Laboratory).

Hamiaux, C., Drummond, R.S.M., Janssen, B.J., Ledger, S.E., Cooney, J.M., Newcomb, R.D., et al. (2012) DAD2 Is an α/β Hydrolase Likely to Be Involved in the Perception of the Plant Branching Hormone, Strigolactone. Current Biology. 22: 2032–2036.

Harris, B.J., Clark, J.W., Schrempf, D., Szöllősi, G.J., Donoghue, P.C.J., Hetherington, A.M., et al. (2022) Divergent evolutionary trajectories of bryophytes and tracheophytes from a complex common ancestor of land plants. Nature Ecology & Evolution. 6: 1634–1643.

Harrison, C.J. (2016) Development and genetics in the evolution of land plant body plans. Philosophical Transactions of the Royal Society B Biological Sciences. 372: 20150490.

Harrison, C.J., and Morris, J.L. (2017) The origin and early evolution of vascular plant shoots and leaves. Philosophical Transactions of the Royal Society B Biological Sciences. 373: 20160496.

Hassan, M.M., Zhang, Y., Yuan, G., De, K., Chen, J.-G., Muchero, W., et al. (2021) Construct design for CRISPR/Cas-based genome editing in plants. Trends in Plant Science. 26: 1133– 1152.

Huang, X., Wang, W., Gong, T., Wickell, D., Kuo, L.-Y., Zhang, X., et al. (2022) The flying spider-monkey tree fern genome provides insights into fern evolution and arborescence. Nature Plants. 8: 500–512.

I, P. (2016) A community-derived classification for extant lycophytes and ferns. Journal of Systematics and Evolution. 54: 563–603.

Jiang, W., Deng, F., Babla, M., Chen, C., Yang, D., Tong, T., et al. (2024) Efficient gene editing of a model fern species through gametophyte-based transformation. PLANT PHYSIOLOGY. 196: 2346–2361.

Kenrick, P., and Crane, P.R. (1997) The origin and early evolution of plants on land. Nature. 389: 33–39.

Kinosian, S.P., and Wolf, P.G. (2022) The biology of C. richardii as a tool to understand plant evolution. eLife. 11.

Kodama, K., Rich, M.K., Yoda, A., Shimazaki, S., Xie, X., Akiyama, K., et al. (2022) An ancestral function of strigolactones as symbiotic rhizosphere signals. Nature Communications. 13: 3974.

Kodama, K., Xie, X., and Kyozuka, J. (2023) The D14 and KAI2 orthologs of gymnosperms sense strigolactones and KL mimics, respectively, and the signals are transduced to control downstream genes. Plant and Cell Physiology. 64: 1057–1065.

Komatsu, A., Fujibayashi, M., Kumagai, K., Suzuki, H., Hata, Y., Takebayashi, Y., et al. (2025) KAI2-dependent signaling controls vegetative reproduction in Marchantia polymorpha through activation of LOG-mediated cytokinin synthesis. Nature Communications. 16: 1263.

Komatsu, A., Kodama, K., Mizuno, Y., Fujibayashi, M., Naramoto, S., and Kyozuka, J. (2023) Control of vegetative reproduction in Marchantia polymorpha by the KAI2-ligand signaling pathway. Current Biology. 33: 1196-1210.e4.

Lai, D.N., Yang, X., Xie, C., Li, T., Yan, A., Liu, X., et al. (2026) Dynamic auxin maxima regulate male-to-hermaphrodite conversion and de novo meristem formation in the fern Ceratopteris gametophytes. PLoS Biology. 24: e3003592.

Li, F.-W., Brouwer, P., Carretero-Paulet, L., Cheng, S., De Vries, J., Delaux, P.-M., et al. (2018) Fern genomes elucidate land plant evolution and cyanobacterial symbioses. Nature Plants. 4: 460–472.

Lopez-Obando, M., Guillory, A., Boyer, F.-D., Cornu, D., Hoffmann, B., Bris, P.L., et al. (2021) The Physcomitrium (Physcomitrella) patens PpKAI2L receptors for strigolactones and related compounds function via MAX2-dependent and -independent pathways. The Plant Cell. 33: 3487–3512.

Marchant, D.B., Chen, G., Cai, S., Chen, F., Schafran, P., Jenkins, J., et al. (2022) Dynamic genome evolution in a model fern. Nature Plants. 8: 1038–1051.

McConnell, H., Lanclos, J.R., Willis, K., Gjording, N., Stockmann, G., Lind, C., et al. (2025) LEAFY demonstrates functions in reproductive development of the gametophyte but not the sporophyte of the fern Ceratopteris richardii. Development. 153.

Mizuno, Y., Komatsu, A., Shimazaki, S., Naramoto, S., Inoue, K., Xie, X., et al. (2021) Major components of the KARRIKIN INSENSITIVE2-dependent signaling pathway are conserved in the liverwort Marchantia polymorpha. The Plant Cell. 33: 2395–2411.

Molla, K.A., Sretenovic, S., Bansal, K.C., and Qi, Y. (2021) Precise plant genome editing using base editors and prime editors. Nature Plants. 7: 1166–1187.

Muthukumar, B., Joyce, B.L., Elless, M.P., and Stewart, C.N. (2013) Stable Transformation of Ferns Using Spores as Targets: Pteris vittata and Ceratopteris thalictroides. PLANT PHYSIOLOGY. 163: 648–658.

Nelson, D.C., Flematti, G.R., Ghisalberti, E.L., Dixon, K.W., and Smith, S.M. (2012) Regulation of Seed Germination and Seedling Growth by Chemical Signals from Burning Vegetation. Annual Review of Plant Biology. 63: 107–130.

Pelosi, J.A., Davenport, R., Kuo, L.-Y., Gray, L.N., Dant, A.J., Kim, E.H., et al. (2025) The genome of the vining fern Lygodium microphyllum highlights genomic and functional differences between life phases of an invasive plant. Proceedings of the National Academy of Sciences. 122: e2504773122.

Plackett, A.R., Conway, S.J., Hazelton, K.D.H., Rabbinowitsch, E.H., Langdale, J.A., and Di Stilio, V.S. (2018) LEAFY maintains apical stem cell activity during shoot development in the fern Ceratopteris richardii. eLife. 7.

Plackett, Andrew R. G., Di Stilio, V.S., and Langdale, J.A. (2015) Ferns: the missing link in shoot evolution and development. Frontiers in Plant Science. 6: 972.

Plackett, A.R.G., Huang, L., Sanders, H.L., and Langdale, J.A. (2014) High-Efficiency Stable Transformation of the Model Fern Species Ceratopteris richardii via Microparticle Bombardment. PLANT PHYSIOLOGY. 165: 3–14.

Plackett, Andrew R G, Rabbinowitsch, E.H., and Langdale, J.A. (2015) Protocol: genetic transformation of the fern Ceratopteris richardii through microparticle bombardment. Plant Methods. 11: 37.

Qin, G., Pan, D., Long, Y., Lan, H., Guan, D., and Song, J. (2024) Chromosome-Scale Genome of the Fern Cibotium barometz Unveils a Genetic Resource of Medicinal Value. Horticulturae. 10: 1191.

Rahmatpour, N., Kuo, L., Kang, J., Herman, E., Lei, L., Li, M., et al. (2022) Analyses of Marsilea vestita genome and transcriptomes do not support widespread intron retention during spermatogenesis. New Phytologist. 237: 1490–1494.

Renninger, K.A., Yarvis, R.M., Youngstrom, C.E., and Cheng, C. (2024) The rise of CLAVATA: evidence for CLAVATA3 and WOX signaling in the fern gametophyte. The Plant Journal. 121: e17207.

Schulz, R., and Theißen, G. (2025) Ceratopteris richardii U6 promoters and plants expressing Cas9 endonuclease as tools for efficient genome editing in a fern. bioRxiv (Cold Spring Harbor Laboratory).

Sepulveda, C., Guzmán, M.A., Li, Q., Villaécija-Aguilar, J.A., Martinez, S.E., Kamran, M., et al. (2022) KARRIKIN UP-REGULATED F-BOX 1 (KUF1) imposes negative feedback regulation of karrikin and KAI2 ligand metabolism in Arabidopsis thaliana. Proceedings of the National Academy of Sciences. 119: e2112820119.

Shu, J., Zhang, Y., Huang, T., and Yan, Y. (2025) The chromosome-level genome assembly of Broad-Leaf Fern (Dipteris shenzhenensis). Scientific Data. 12: 475.

Shukla, A.K., Upadhyay, S.K., Mishra, M., Saurabh, S., Singh, R., Singh, H., et al. (2016) Expression of an insecticidal fern protein in cotton protects against whitefly. Nature Biotechnology. 34: 1046–1051.

Song, M.J., Rizzieri, Y.C., Li, F.-W., Freund, F., Escalona, M., Toffelmier, E., et al. (2025) The genome assembly of the duckweed fern, Azolla caroliniana. Journal of Heredity. 116: 691–701.

Stanga, J.P., Morffy, N., and Nelson, D.C. (2016) Functional redundancy in the control of seedling growth by the karrikin signaling pathway. Planta. 243: 1397–1406.

Stanga, J.P., Smith, S.M., Briggs, W.R., and Nelson, D.C. (2013) SUPPRESSOR OF MORE AXILLARY GROWTH2 1 controls seed germination and seedling development in Arabidopsis. PLANT PHYSIOLOGY. 163: 318–330.

Stuttmann, J., Barthel, K., Martin, P., Ordon, J., Erickson, J.L., Herr, R., et al. (2021) Highly efficient multiplex editing: one-shot generation of 8× Nicotiana benthamiana and 12× Arabidopsis mutants. The Plant Journal. 106: 8–22.

Sugano, S.S., Nishihama, R., Shirakawa, M., Takagi, J., Matsuda, Y., Ishida, S., et al. (2018) Efficient CRISPR/Cas9-based genome editing and its application to conditional genetic analysis in Marchantia polymorpha. PLoS ONE. 13: e0205117.

Tuncel, A., Pan, C., Clem, J.S., Liu, D., and Qi, Y. (2025) CRISPR–Cas applications in agriculture and plant research. Nature Reviews Molecular Cell Biology. 26: 419–441.

Wang, L., Wang, B., Jiang, L., Liu, X., Li, X., Lu, Z., et al. (2015) Strigolactone signaling in Arabidopsis regulates shoot development by targeting D53-Like SMXL repressor proteins for ubiquitination and degradation. The Plant Cell. 27: 3128–3142.

Waters, M.T., and Smith, S.M. (2012) KAI2- and MAX2-Mediated responses to karrikins and strigolactones are largely independent of HY5 in arabidopsis seedlings. Molecular Plant. 6: 63– 75.

Withers, K.A., Falls, K., Youngstrom, C.E., Nguyen, T., DeWald, A., Yarvis, R.M., et al. (2023) A Ceratopteris EXCESS MICROSPOROCYTES1 suppresses reproductive transition in the fern vegetative leaves. Plant Science. 335: 111812.

Xia, Z., Duan, L., Fang, Y., Jiang, Y., Chen, H., Yan, Y., et al. (2025) Decoding the genome of Brainea insignis reveals insights into fern evolution and conservation. Nature Communications. 17: 1292.

Xiang, D.-L., and Li, G.-S. (2024) Control of leaf development in the water fern Ceratopteris richardii by the auxin efflux transporter CrPINMa in the CRISPR/Cas9 analysis. BMC Plant Biology. 24: 322.

Yang, X., Yan, A., Liu, X., Volkening, A., and Zhou, Y. (2025) Single cell-derived multicellular meristem: insights into male-to-hermaphrodite conversion and de novo meristem formation in Ceratopteris. Development. 152.

Yang, Y., Yang, Yanqiu, Deng, S., and Ying, Z. (2025) Role of Azolla in sustainable agriculture and climate resilience: a comprehensive review. Frontiers in Plant Science. 16: 1661720.

Youngstrom, C.E., Geadelmann, L.F., Irish, E.E., and Cheng, C.-L. (2019) A fern WUSCHEL-RELATED HOMEOBOX gene functions in both gametophyte and sporophyte generations. BMC Plant Biology. 19: 416.

Youngstrom, C.E., Withers, K.A., Irish, E.E., and Cheng, C.-L. (2022) Vascular function of the T3/modern clade WUSCHEL-Related HOMEOBOX transcription factor genes predate apical meristem-maintenance function. BMC Plant Biology. 22: 210.

Zhang, Y., Werling, U., and Edelmann, W. (2012) SLiCE: a novel bacterial cell extract-based DNA cloning method. Nucleic Acids Research. 40: e55.

Zhang, Yingxiao, Ren, Q., Tang, X., Liu, S., Malzahn, A.A., Zhou, J., et al. (2021) Expanding the scope of plant genome engineering with Cas12a orthologs and highly multiplexable editing systems. Nature Communications. 12: 1944.

Zhong, Y., Liu, Y., Wu, W., Chen, J., Sun, C., Liu, H., et al. (2022) Genomic Insights into Genetic Diploidization in the Homosporous Fern Adiantum nelumboides. Genome Biology and Evolution. 14.

