## Supplementary Figure S1 for "Generation of *KARRIKIN INSENSITIVE2* loss-of-function mutants in *Ceratopteris richardii* using a CRISPR/Cas9 system based on ribozyme-gRNA-ribozyme (RGR) technology"

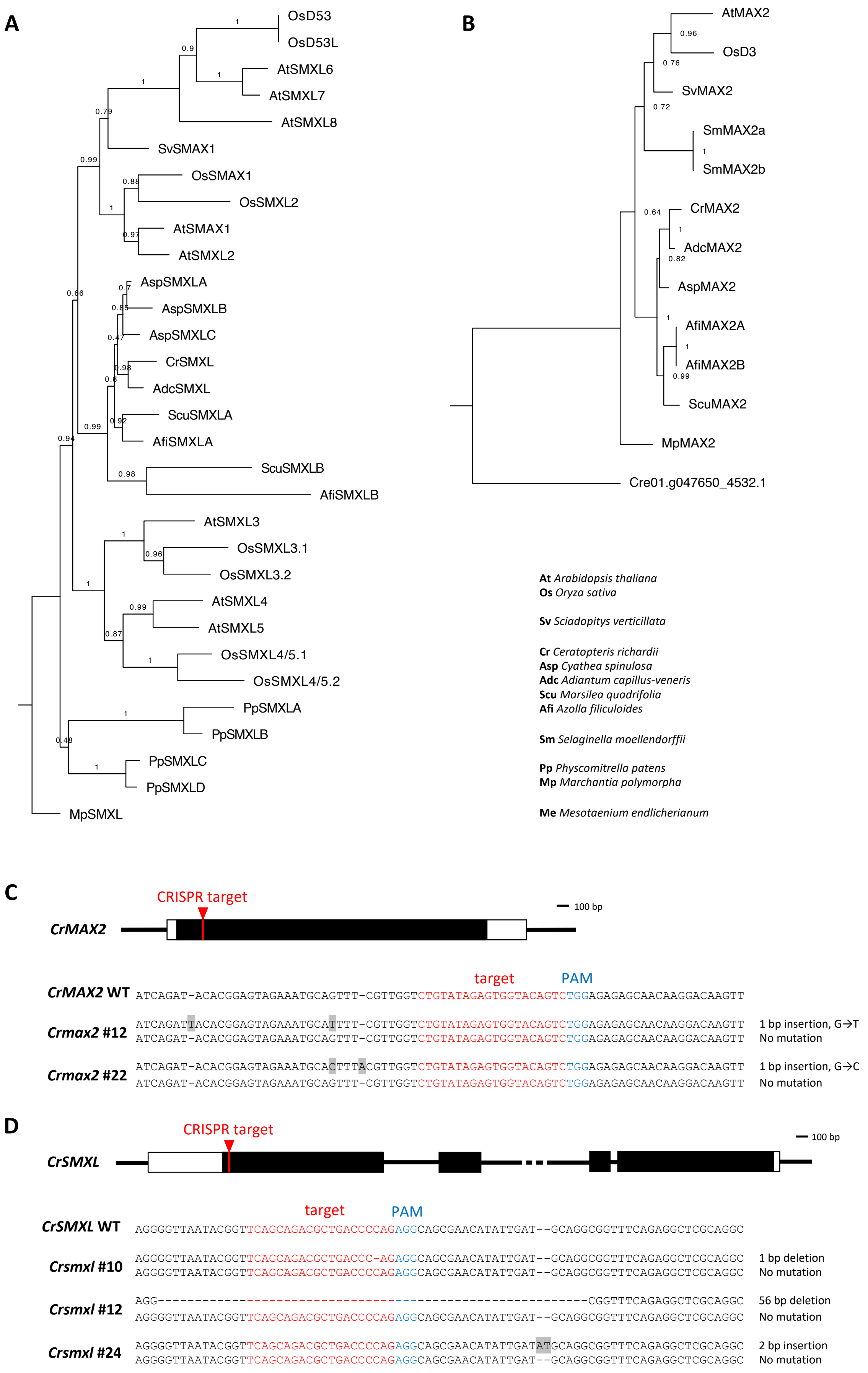


**Figure S1 Generation of *Crsmxl* and *Crmax2* loss-of-function mutants**
**(A and B)** Phylogenetic tree of SMXL family (A) and MAX2 family (B) proteins.
**(C and D)** Gene structure of *CrSMXL* (C) and *CrMAX2* (D). Introduced mutation patterns at the CRISPR/Cas9 target site are shown below. White and black boxes indicate exon and protein-coding sequences, respectively.
