## Supplementary Table S1 for "Generation of *KARRIKIN INSENSITIVE2* loss-of-function mutants in *Ceratopteris richardii* using a CRISPR/Cas9 system based on ribozyme-gRNA-ribozyme (RGR) technology"

### Table S1 Oligos used in this study

| No. | name | sequence (5'→3') |
|---|---|---|
| #1 | CrKAI2_68fw_OligoA | CTCGAGGGAACGTGGTGGTAGTCC |
| #2 | CrKAI2_68fw_OligoB | AAACGGACTACCACCACGTTCCCT |
| #3 | 35S_ABP1_RGR_F_CACC | CACCAACATGGTGGAGCACGACAC |
| #4 | 35S_ABP1_RGR_R | GTTTGGGGATCTAGTGTTTTACTCC |
| #5 | RGR_CrKAI2_F1 | CAGCCTCTACACGTTTAAACTTCCCTCTGATGAGTCCGTGAGGACGAAACGAGTAAGCTCGTC |
| #6 | RGR_CrKAI2_F2 | GACGAAACGAGTAAGCTCGTCAGGGAACGTGGTGGTAGTCCGTTTTAGAGCTAGAAATAGCAAG |
| #7 | RGR_CrMAX2_F1 | CAGCCTCTACACGTTTAAACATACAGCTGATGAGTCCGTGAGGACGAAACGAGTAAGCTCGTC |
| #8 | RGR_CrMAX2_F2 | GACGAAACGAGTAAGCTCGTCCTGTATAGAGTGGTACAGTCGTTTTAGAGCTAGAAATAGCAAG |
| #9 | RGR_CrSMXL_F1 | CAGCCTCTACACGTTTAAACTGCTGACTGATGAGTCCGTGAGGACGAAACGAGTAAGCTCGTC |
| #10 | RGR_CrSMXL_F2 | GACGAAACGAGTAAGCTCGTCTCAGCAGACGCTGACCCCAGGTTTTAGAGCTAGAAATAGCAAG |
| #11 | RGR_R | AGTTAGGTCTAGACTAGTTTGTCCCATTCGCCATGCCGAAGC |
| #12 | Cas9_Pea3Ter_F_CACC | CACCATGGATAAGAAGTACTCTATCGG |
| #13 | Cas9_Pea3Ter_R | AAGCCTATACTGTACTTAACTTGATT |
| #14 | SLiCE_Gateway_F | GTGGTTGATAACAGCGGTTGACTAGAGTTATCAACAAGTT |
| #15 | SLiCE_Gateway_R | ATTCGAGCTCTAAGCATTCGAGCTCTAAGCGCTGT |
| #16 | Hyg91F | GGCGAAGAATCTCGTGCTTTC |
| #17 | Hyg696R | GATGTTGGCGACCTCGTATT |
| #18 | CrKAI2_Crispr_check_F | CCAGCAATTTTCATGGAGCGT |
| #19 | CrKAI2_Crispr_check_R | CTGGGAGACGCAGAGACAAG |
| #20 | CrMAX2_Crispr_check_F | TCCCGCTTCCATGTAAATCAGA |
| #21 | CrMAX2_Crispr_check_R | GGTCAAGGGACTCCACTGC |
| #22 | CrSMXL_Crispr_check_F | TAGTCGGCACACTTCCTTGC |
| #23 | CrSMXL_Crispr_check_R | CACGTTTGAGTGCAGCAACA |
| #24 | qCrKUF_F | GCTTTACGCCCTCCTTCACG |
| #25 | qCrKUF_R | AGCGCATGCCATGAGTTAGC |
| #26 | qCrDDK_F | CTTGCAGCACTGACCGAACC |
| #27 | qCrDDK_R | ACCACGATGTTTGCAGTGCC |
| #28 | qCrSMXL_F | CTGCAGAAGGCTCTTCCTGC |
| #29 | qCrSMXL_R | CGCAACTCTTGTAACCTCATGC |
| #30 | qCrTBPb_F | ATGAGCCAGAGCTTTTCCCC |
| #31 | qCrTBPb_R | TTCGTCTCTGACCTTTGCCC |
